# Single-cell RNA-sequencing identifies unique cell-specific gene expression profiles in high-grade cardiac allograft vasculopathy

**DOI:** 10.1101/2024.07.10.602989

**Authors:** Kaushik Amancherla, Kelly H. Schlendorf, Nelson Chow, Quanhu Sheng, Jane E. Freedman, Jeffrey C. Rathmell

**Author notes:** **Corresponding author:** Kaushik Amancherla, MD, Instructor in Medicine, Vanderbilt University Medical Center, Nashville, TN. These authors contributed equally as senior authors (Jane E. Freedman, Jeffrey C. Rathmell).

## Abstract

**Background:** Cardiac allograft vasculopathy (CAV), a diffuse thickening of the intima of the coronary arteries and microvasculature, is the leading cause of late graft failure and mortality after heart transplantation (HT). Diagnosis involves invasive coronary angiography, which carries substantial risk, and minimally-invasive approaches to CAV diagnosis are urgently needed. Using single-cell RNA-sequencing in peripheral blood mononuclear cells (PBMCs), we sought to identify cell-specific gene expression profiles in CAV.

**Methods:** Whole blood was collected from 22 HT recipients with angiographically-confirmed CAV and 18 HT recipients without CAV. PBMCs were isolated and subjected to single-cell RNA-sequencing using a 10X Genomics microfluidic platform. Downstream analyses focused on differential expression of genes, cell compositional changes, and T cell receptor repertoire analyses.

**Results:** Across 40 PBMC samples, we isolated 134,984 cells spanning 8 major clusters and 31 subclusters of cell types. Compositional analyses showed subtle, but significant increases in CD4+ T central memory cells, and CD14+ and CD16+ monocytes in high-grade CAV (CAV-2 and CAV-3) as compared to low-grade or absent CAV. After adjusting for age, gender, and prednisone use, 745 genes were differentially expressed in a cell-specific manner in high-grade CAV. Weighted gene co-expression network analyses showed enrichment for putative pathways involved in inflammation and angiogenesis. There were no significant differences in T cell clonality or diversity with increasing CAV severity.

**Conclusions:** Unbiased whole transcriptomic analyses at single-cell resolution identify unique, cell-specific gene expression patterns in CAV, suggesting the potential utility of peripheral gene expression biomarkers in diagnosing CAV.

## Introduction

Heart transplantation (HT) is the definitive treatment for end-stage heart failure. However, despite substantial improvements in early post-HT survival in recent decades, graft failure due to cardiac allograft vasculopathy (CAV) remains a leading cause of death beyond the first-year post-HT^1^. Coronary angiography, performed with adjunctive intravascular ultrasound at some transplant centers, remains the gold standard for CAV diagnosis^2^. However, this procedure is invasive (carrying risk of complications, including life-threatening ones^3, 4^), requires use of nephrotoxic contrast agents, and is costly^5^. Moreover, coronary angiography often fails to detect CAV until late in its course. Novel non-invasive biomarkers for earlier CAV diagnosis may overcome these barriers and have the potential to substantially impact patient care.

Peripheral biomarkers have been developed for acute rejection but have yet to show significant clinical utility in CAV^6–17^. Gene expression profiling (e.g., *Allomap*) of peripheral blood mononuclear cells (PBMCs) using targeted gene panels has been developed for non-invasive detection of acute cellular rejection and is currently commercially available^6–9^. More recently, donor-derived cell-free DNA (dd-cfDNA) has shown promise in early detection of acute rejection^18^. However, targeted peripheral gene panels have not shown consistent association with CAV and there are limited data regarding dd-cfDNA in CAV^19, 20^. Small studies assessing targeted proteomics^15–17, 21^, serum cytokine levels^12^, and serum levels of microRNAs (miRNAs)^10,11^ in CAV have been proposed but show limited sensitivity and specificity.

Single-cell RNA-sequencing (scRNA-seq) has emerged as a powerful tool to characterize the whole transcriptome, rather than targeting specific genes, at the level of individual cells. This approach can identify rare cell types/states that are perturbed in diseased conditions and characterize dynamic cell-specific transcriptomic profiles. We have previously defined cell-specific heterogeneity and donor-recipient chimerism in human cardiac biopsies in severe CAV^22^. In this study, we leverage scRNA-seq capabilities to explore cell-specific gene expression profiles and T cells receptor (TCR) repertoires in PBMCs of patients with angiographically-confirmed CAV.

## Methods

### Human samples

The overall study design is depicted in **Figure 1A**. Adult HT recipients were enrolled during routine transplant clinic visits at Vanderbilt University Medical Center. Eligible participants were screened for presence or absence of CAV and approached for study participation in May 2022. All participants provided written informed consent. This study was approved by the Vanderbilt University Institutional Review Board (#200551). Peripheral blood (∼20 mL) was collected in vacutainers containing K2 EDTA, sodium heparin, or acid citric dextrose solution A.

**Figure 1.**
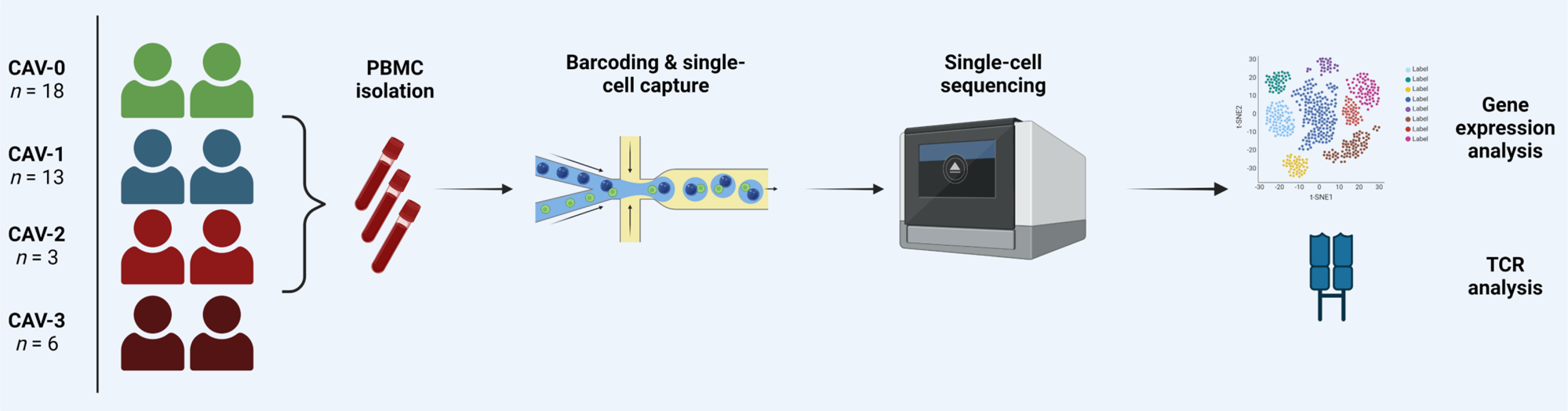
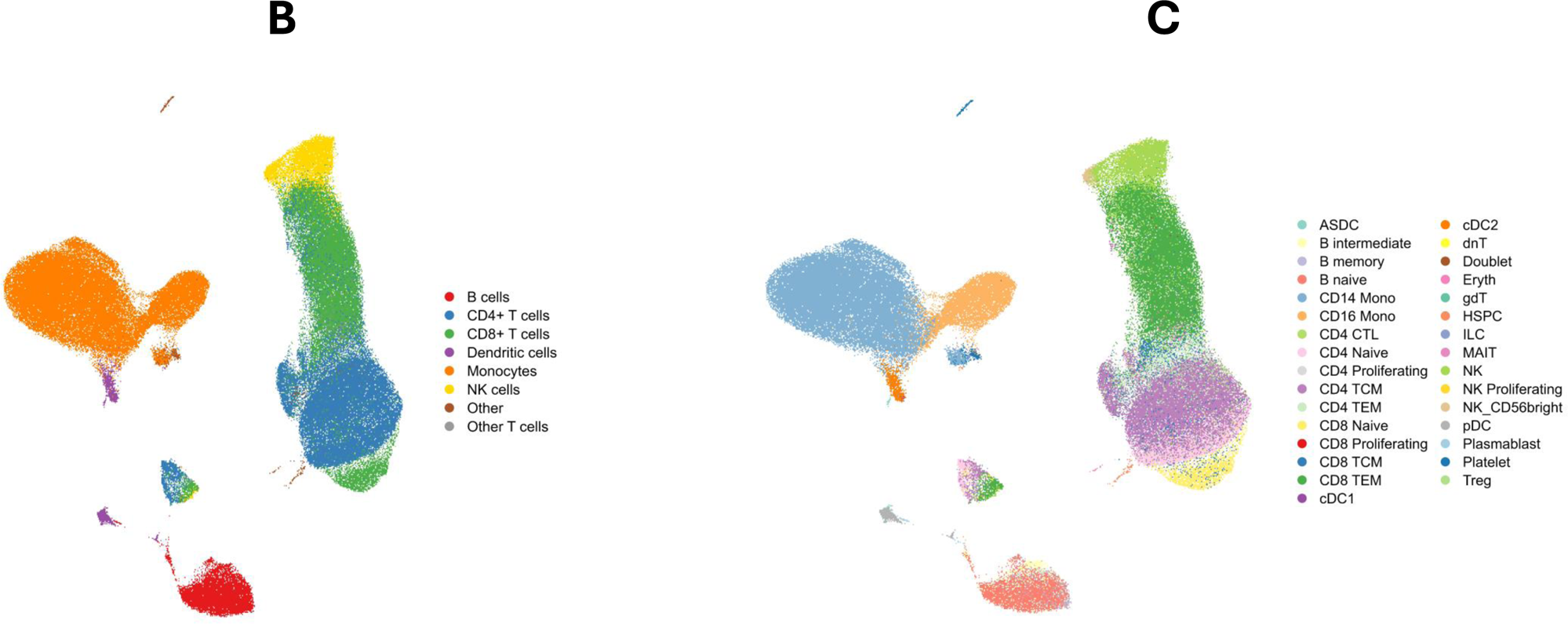
**A)** Overview of the study design and samples collected. This figure was created using Biorender.com. **B)** UMAP visualization of 134,984 cells passing quality control, as described. Using canonical markers and reference-mapping, 8 major clusters were defined. **C)** 31 subclusters were further characterized. ASDC = *AXL+ SIGLEC6+* dendritic cells; Mono = monocyte; CTL = cytotoxic T lymphocytes; TCM = T central memory cells; TEM = T effector memory cells; cDC = conventional dendritic cells; dnT = double negative T cells; gdT = γδ T cells; HSPC = hematopoietic stem and progenitor cells; ILC = circulating innate lymphoid cells; MAIT = mucosal-associated invariant T cells; NK = natural killer cells; pDC = plasmacytoid dendritic cells; Treg = T regulatory cells.

### PBMC isolation

Briefly, whole blood was diluted with PBS (Mediatech 21-031-CM). The blood/PBS mixture was layered onto Ficoll-Paque (GE life science Cat# 17-1440-03) and centrifuged at 400g for 30 minutes at room temperature. Following centrifugation, the supernatant was removed by aspiration, the PBMC layer was collected, and cells were washed twice with PBS. Red blood cells were removed through incubation for 5 minutes at room temperature in ACK Lysis Solution (Gibco Cat# A 10492-01). Cells were counted and underwent a final wash, followed by gentle addition of freezing medium (10% DMSO/90% FBS; Sigma D2650 and Atlas Biologicals Cat# F-0500-A) and aliquoting into vials. Vials were placed in −80C overnight and transferred to a liquid nitrogen freezer the following day.

### Single-cell RNA-sequencing

Prior to scRNA-seq assays, cryopreserved vials were thawed rapidly (∼30 seconds) in a 37C water bath. RPMI (LifeTechnologies, MT15040CV) was added and vials were centrifuged at 400g for 7 minutes. Supernatant was removed and the cell pellet was resuspended. Following another round of centrifugation at 400g for 7 minutes, cells were counted and resuspended. Next, 5 µL of Human TruStain FcX Fc Blocking Reagent was added to the cells, followed by incubation for 10 minutes on ice, addition of 1 µL of TotalSeq C hashtag antibodies, and re-incubation for 30 minutes on ice. Cells were resuspended in 5 mL PBS and 1% BSA, centrifuged at 400g for 10 minutes, and a final suspension was brought to ∼3000 cells/µL.

Samples labeled with TotalSeq C hashtag antibodies were pooled together to generate mixtures of 5 samples per batch (total 8 batches). Pooled samples were loaded into a 10X Genomics microfluidic platform (Single Cell 5’ v2 Reagent Kit), targeting 12,000 – 16,000 cells per batch. Sequencing was performed on the NovaSeq6000 S4 flow cell targeting 50,000 reads per cell for the gene expression libraries and 5,000 reads per cell for the feature barcode libraries. Libraries were processed according to manufacturer’s instructions.

The raw FASTQ files were aligned to the human genome (GRCh38) and an RNA count matrix was generated using CellRanger (v6.1.2). Count matrices were imported into R (v4.2.2) and postprocessing was performed using Seurat (v4.3)^23^. Ambient RNA was corrected for using DecontX (v1.0.0) at the batch level^24^. Cells with nFeature_RNA and nCount_RNA greater than the 75^th^ percentile + 1.5x the interquartile range (IQR) were removed from downstream analyses. Doublets were filtered using scDblFinder (v1.12.0)^25^ and cells with more than 10% of genes mapped to mitochondrial genes were removed. Batches were demultiplexed using the *HTODemux* function (Seurat v4.3).

TCR and immunoglobulin genes, which are highly variable, were removed to minimize their influence on downstream clustering. Each batch underwent normalization using scTransform v2 (v0.3.5)^26^. Principal components (PCs) were calculated using 3,000 highly variable genes and corrected for batch effects using Harmony (v1.1)^27^. Using batch-corrected PCs, the nearest neighbors were calculated to generate a uniform manifold approximation and projection (UMAP). Clustering was performed using the Louvain algorithm at a resolution of 0.4. Cell type identification was performed using a combination of marker gene identification and reference mapping to a published cellular indexing of transcriptomes and epitopes by sequencing (CITE-seq) data set of PBMCs^28^.

We focused our comparisons on high-grade CAV (CAV-2 or CAV-3) versus low-grade CAV (CAV-0 and CAV-1) due to recent data suggesting that patients with CAV-0 and CAV-1 have similar long-term risks of graft failure distinct from those with higher grades of CAV^29, 30^. Differences in cell composition were performed using Milo (v1.6.0)^31^. Milo tests for differential abundance within neighborhoods on the k-nearest neighbor (kNN) graph using a negative binomial generalized linear model. This approach does not rely on discrete clustering of cell types, allowing for the identification of subtle changes in neighborhoods within globally-annotated clusters along the continuous spectrum of PBMC differentiation. We constructed the kNN graph using *k* = 90, to target ∼200-250 cells per neighborhood, and used 30 PCs (*d* = 30). This approach defined 9,371 neighborhoods spread across all clusters. Differences in numbers of cells across samples within each neighborhood was normalized by trimmed mean of M-values. Differential abundance was tested in each neighborhood using the quasi-likelihood method in edgeR^32^. Multiple testing correction is performed on nominal P-values through a weighted Benjamini-Hochberg method, as previously described. Neighborhoods were considered differentially abundant if they met a spatial false discovery rate (FDR)-adjusted P-value < 0.1.

As immune cells exist on a continuous spectrum of differentiation, we used miloDE (v0.0.0.9)^33^ – a cluster-free differential expression (DE) analysis framework – to increase sensitivity for DE gene detection. MiloDE assigns cells with homogenous transcriptomic profiles into neighborhoods using a 2^nd^ order kNN graph on the batch-corrected Harmony embedding. We targeted a median of ∼1,000 cells per neighborhood and DE analyses were conducted within each neighborhood. After initial neighborhood assignment, we excluded neighborhoods that were unlikely to show any DE genes through utilization of a random forest classifier (excluding neighborhoods with area under the curve [AUC] ≤ 0.5), leaving 424 neighborhoods for DE testing and minimizing the penalty of multiple testing correction. Conditions were defined as 1) high-grade CAV (grouping CAV-2 and CAV-3 together; as compared with CAV-0 and CAV-1) and 2) CAV (grouping CAV-1, CAV-2, and CAV-3 together; as compared with CAV-0). DE analyses included age, gender, and prednisone use as covariates. Multiple testing correction was performed using the Benjamini-Hochberg method for all tested genes within each neighborhood and across neighborhoods. Genes were considered significantly DE if they met an FDR-adjusted P-value < 0.05. To identify co-regulated transcriptional programs, significantly DE genes in at least 2-4 neighborhoods were input into scWGCNA (v1.0)^34^ at an exploratory P-value threshold of 0.1 to detect gene modules. Pathway enrichment analyses, at both the pseudobulk (all DE genes within neighborhoods) and module levels, were performed using clusterProfiler (v4.6.2) with all tested genes as the background universe^35^. The org.Hs.eg.db (v3.16.0) package was used for genome annotation and DOSE (v3.24.2)^36^ was used for ontology terms, with Biological Process (BP) terms prioritized. Multiple testing correction was performed using the Benjamini-Hochberg method.

### Single-cell T cell receptor analyses

BAM files generated after alignment with the human genome (GRCh38) were used to infer CDR3 sequences of TCRs using TRUST4 (v1.0.13)^37^. Annotation and quantification of TCR clonotypes was performed using scRepertoire (v2.0.3)^38^. The barcodes for each clone were intersected with barcodes from scRNA-seq at a batch level to filter down to high-quality T cells that passed scRNA-seq quality metrics, as described above. Nonproductive chains were removed. Proportional cut points for clonal frequencies were 1) rare (0 < X ≤ 0.0001), 2) small (0.0001 < X ≤ 0.001), 3) medium (0.001 < X ≤ 0.01), 4) large (0.01 < X ≤ 0.1), and 5) hyperexpanded (0.1 < X ≤ 1.0). Clonal diversity metrics included the Shannon, inverse Simpson, normalized entropy, Gini-Simpson, Chao1, and abundance-based coverage estimator (ACE) indices. The Kruskal-Wallis test was used to compare repertoire statistics across the four CAV groups, with a P-value < 0.05 considered statistically significant.

### Data and code availability

To ensure scientific rigor and reproducibility, complete code used for analyses is available online at https://github.com/learning-MD/PBMC_CAV. Raw FASTQ files and the processed Seurat object are deposited at the National Institutes of Health/National Center for Biotechnology Gene Expression Omnibus data repository.

## Results

A total of 40 HT recipients were prospectively enrolled in this study, with PBMC samples obtained a median of 61 months post-HT. Of these, 22 patients had angiographically-confirmed CAV: 13 with CAV-1, 3 with CAV-2, and 6 with CAV-3. There was no significant difference in median time from transplant between the CAV and non-CAV groups (71 ± 125 months in the CAV group vs 48.5 ± 55.2 months in the non-CAV group; P-value = 0.10). Additionally, there were no significant differences in major clinical characteristics between the two groups (**Table 1**). Detailed clinical characteristics are provided in **Supplemental Table 1**. Following single-cell isolation and sequencing, 134,984 cells passed quality control. We defined 8 major clusters and 31 subclusters (**Figure 1B**), using a combination of canonical marker genes and reference-mapping with a published CITE-seq study^28^ that quantified cell surface proteins for deep immune phenotyping (**Figure 1C**).

**Table 1.** Clinical characteristics of heart transplant recipients.

| **Demographics by CAV status** | **N** | **No CAV (N = 18)<sup>1</sup>** | **Has CAV (N = 22)<sup>1</sup>** | **p-value<sup>2</sup>** |
| --- | --- | --- | --- | --- |
| **Age (years)** | 40 | 53 (49, 66) | 53 (46, 63) | 0.7 |
| **Gender(% female)** | 40 | 8 (44%) | 10 (45%) | >0.9 |
| **Diabetes mellitus** | 40 | 6 (33%) | 6 (27%) | 0.7 |
| **Time since transplant (months)** | 40 | 49 (23, 78) | 71 (29, 154) | 0.10 |
| **CAV grade** | 40 |  |  | <0.001 |
| 0 |  | 18 (100%) | 0 (0%) |  |
| 1 |  | 0 (0%) | 13 (59%) |  |
| 2 |  | 0 (0%) | 3 (14%) |  |
| 3 |  | 0 (0%) | 6 (27%) |  |
| **Class II DSAs** | 40 | 3 (17%) | 7 (32%) | 0.5 |
| **History of ACR** | 40 | 3 (17%) | 9 (41%) | 0.10 |
| **History of AMR** | 40 | 3 (17%) | 5 (23%) | 0.7 |
| **Prednisone use** | 40 | 5 (28%) | 6 (27%) | >0.9 |
| **Calcineurin inhibitor use** | 40 | 16 (89%) | 19 (86%) | >0.9 |
| **Antimetabolite use** | 40 | 14 (78%) | 16 (73%) | >0.9 |
| **mTOR inhibitor use** | 40 | 5 (28%) | 11 (50%) | 0.2 |

<sup>1</sup>Median (IQR); n (%)
<sup>2</sup>Wilcoxon rank sum test; Pearson's Chi-squared test; Fisher's exact test

We identified 79 differentially abundant neighborhoods at an FDR < 0.1. These subtle changes in cell composition included statistically significant increases in CD4+ T central memory cells (CD4+ TCM) and CD14+ and CD16+ monocytes in individuals with high-grade as compared to low grade CAV (**Figure 2A**). Detailed characteristics of all tested neighborhoods are provided in **Supplemental Table 2**. Comparing high-grade CAV with low-grade CAV, we identified 745 unique genes that were differentially expressed at an FDR < 0.05 (**Figure 2B**). These included genes critical for cytotoxic degranulation (*NKG7*, *GNLY*)^39, 40^, interferon-responsiveness (*IFI27*, *IFI44L*, *IFITM3*, *IFIT1*, *IFIT2*, *IFIT3*, *STAT1*, *CXCL10*)^41^, and angiogenesis (*VEGFA*, *VEGFC*, *FLT1*, *CXCL2*)^42^, among others. In comparing all CAV (CAV-1, CAV-2, and CAV-3) against CAV-0, fewer genes were differentially expressed (250 genes, FDR < 0.05) between groups. The full list of genes, their log2FC, and P-values are in **Supplemental Tables 3 and 4**.

**Figure 2.**
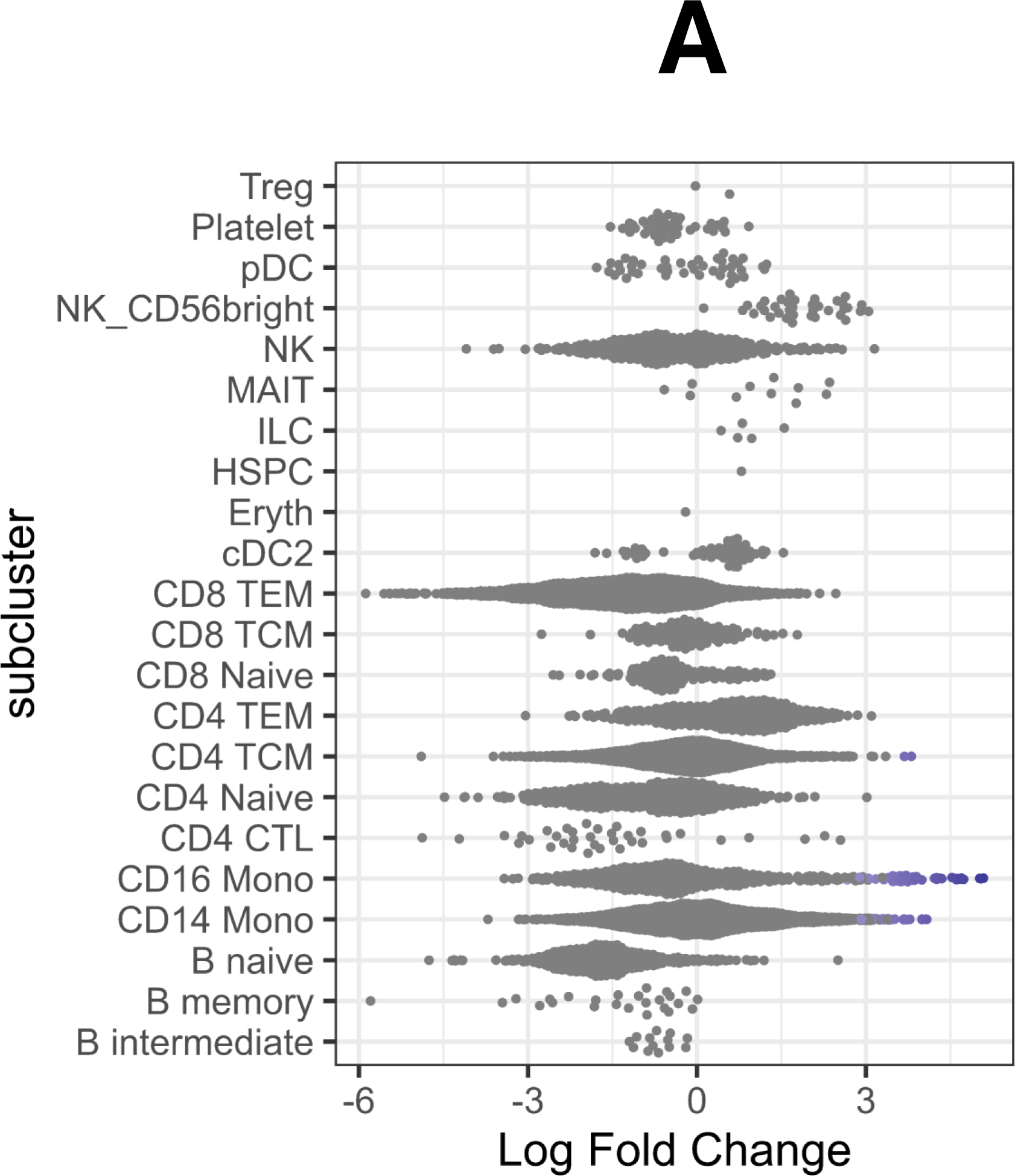
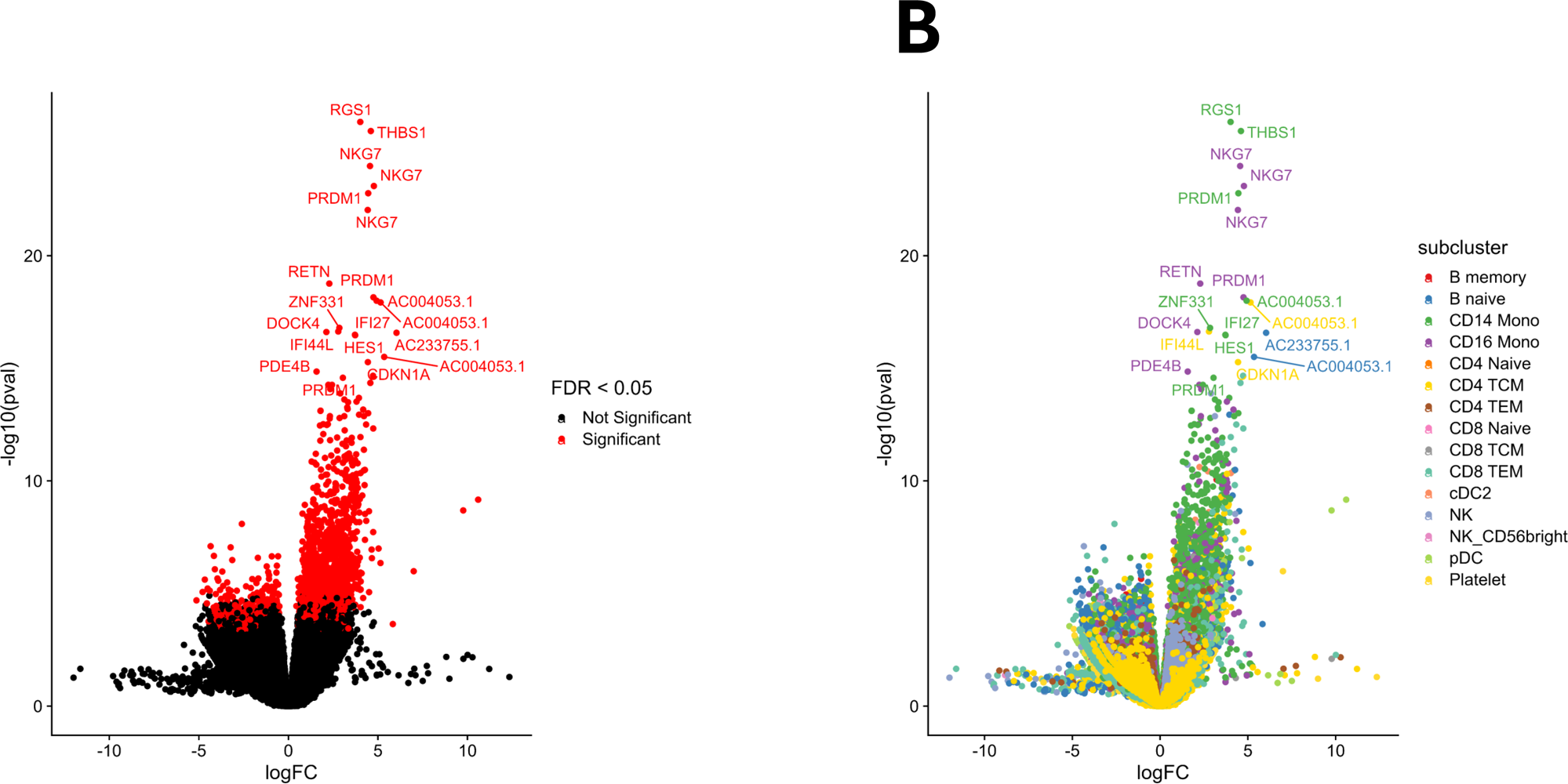
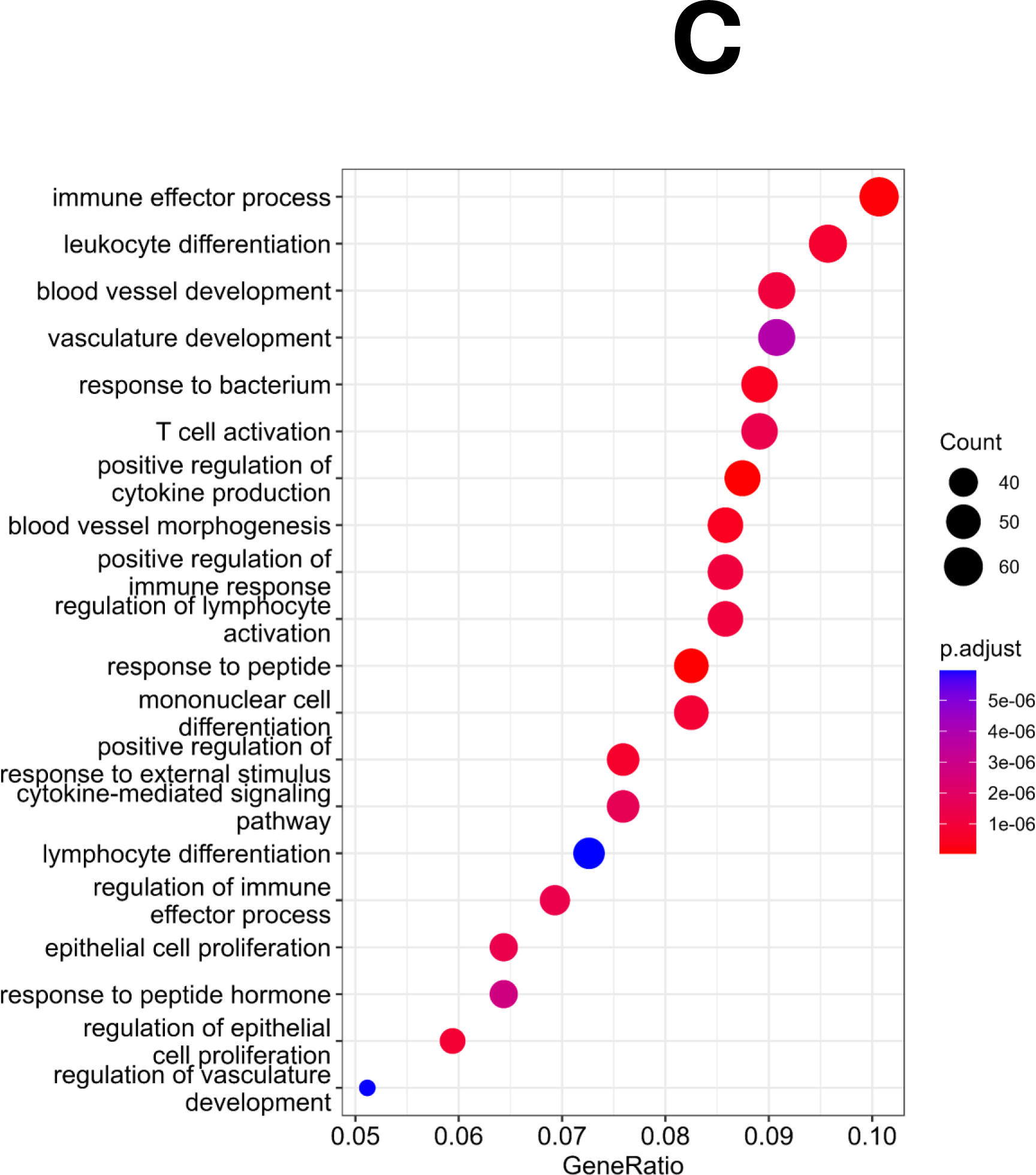
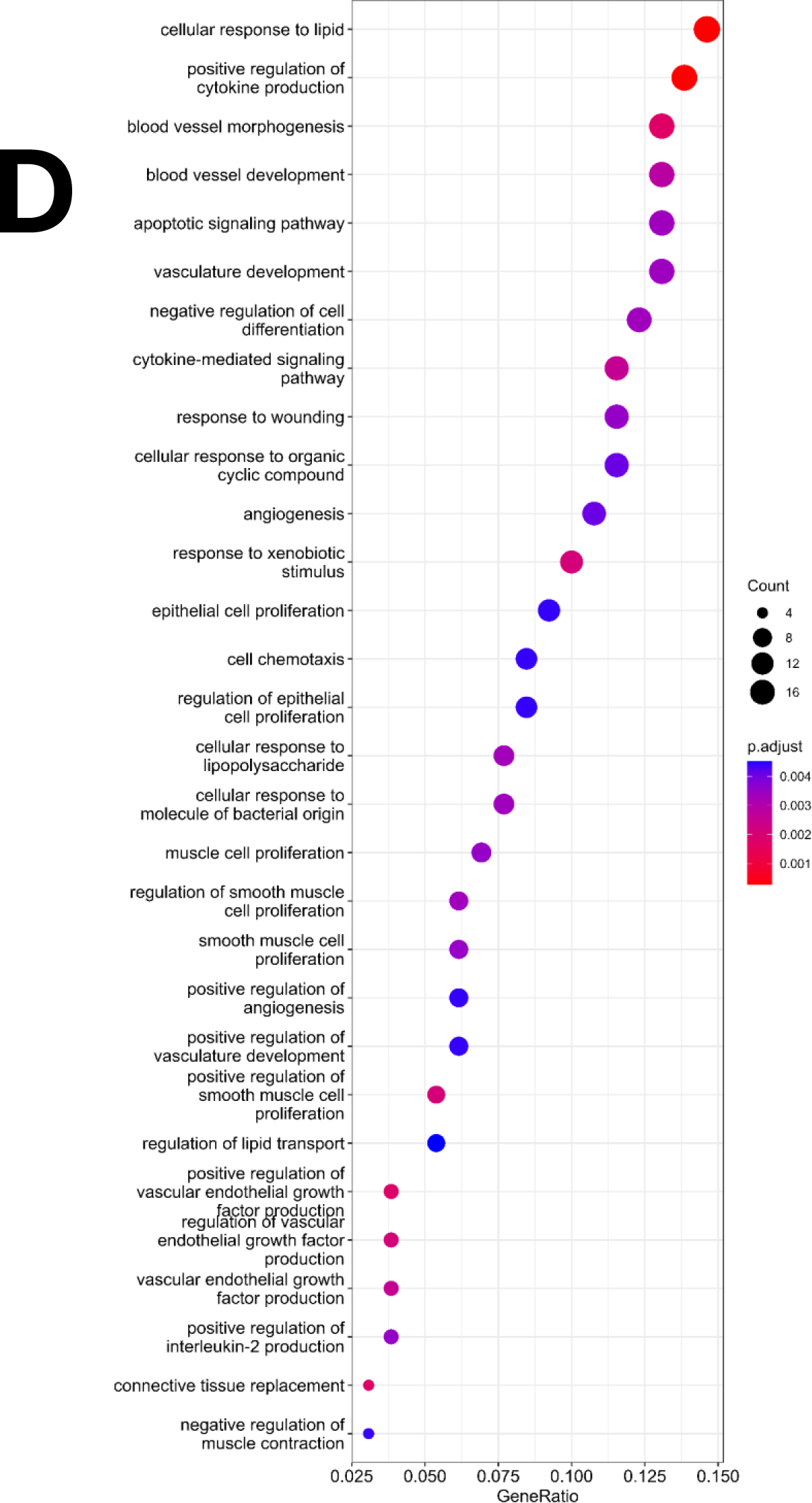
**A)** Beeswarm plot of the distribution of log fold change between high-grade CAV (CAV-2 and CAV-3) and low-grade CAV (CAV-0 and CAV-1) according to neighborhoods within annotated subclusters. Neighborhoods are considered differentially abundant between conditions at an FDR < 0.1 and are colored blue if enriched in high-grade CAV and red if enriched in low-grade CAV. **B)** <u>Left</u>: Volcano plot of differentially expressed genes between high-grade CAV and low-grade CAV. Positive logFC indicates higher expression in high-grade CAV while negative logFC indicates higher expression in low-grade CAV. Red color indicates genes that meet FDR < 0.05 after correction both within tested neighborhoods and across tested neighborhoods. <u>Right</u>: Volcano plot of differentially expressed genes between high-grade CAV and low-grade CAV. Positive logFC indicates higher expression in high-grade CAV while negative logFC indicates higher expression in low-grade CAV. Color of dots indicates the subcluster of the neighborhood within which each gene is differentially expressed. **C)** Using all 745 differentially expressed genes in high-grade CAV, pathway enrichment analysis was performed for Biological Process terms using clusterProfiler (v4.6.2) with all tested genes as the background universe. **D)** Module 7 in high-grade CAV, consists of 156 genes identified via single-cell WGCNA. These 156 genes were tested for pathway enrichment for Biological Process terms using all tested genes as the background universe.

Neighborhoods enriched for CD14+ and CD16+ monocytes highly expressed genes associated with cytotoxic degranulation (*NKG7*^40^, *GNLY*^39^), chemokines and cytokines (e.g., *CCL3L1*^43^, *CCL4L2*^44^, *IL1A*^44, 45^, *CXCL1*^46^, *CXCL2*^42^, *CXCR4*^47^, *CXCL10*^48^, and *IL32*^49^, among others), and interferon-responsiveness (as described above). Both CD4+ and CD8+ T cells similarly expressed interferon-responsive genes and positive regulators of interferon-gamma (e.g., *IFNG-AS1*^50^ in CD8+ T effector memory [TEM] cells). Furthermore, CD4+ TCMs exhibited markers of cytotoxicity (*GZMH*^51^) while CD8+ TEMs cells expressed markers of activation, expansion, and cytotoxicity (*HLA-DRA*, *TNFRSF18*, *CFH*, *STAT1*, etc.)^51^. Neighborhoods enriched for NK cells and dendritic cells highly expressed genes associated with inflammation (e.g., *PRDM1*^52^), interferons and interferon-responsiveness (*IFNG*, *IFNG-AS1*, among others outlined above), and cell migration (e.g., *RUFY3*, *SELL*, etc.)^51, 53^.

Pathway enrichment analysis using all 745 unique DE genes in high-grade CAV showed enrichment for putative pathways in leukocyte differentiation, angiogenesis and the inflammatory response (**Figure 2C**). Subsequent single-cell WGCNA identified 8 modules of co-expressed genes in high-grade CAV of which 5 showed pathway enrichment in a cell-specific manner for biologically-plausible pathways (**Supplemental Figure 1**). Notably, module 4 (differentially-enriched across cell types) showed significant enrichment for pathways involved in immune responses to viral infection (**Supplemental Figure 2A**) and module 7 (consisting of 156 genes; most enriched in CD14+ and CD16+ monocytes) showed significant enrichment for pathways involved in inflammation, angiogenesis, vascular smooth muscle cell proliferation and wound healing (**Figure 2D**). Detailed lists of gene modules for the high-grade vs low-grade CAV and for CAV vs non-CAV comparisons are provided in **Supplemental Table 5 and 6**. Figures of module-specific pathway enrichment for high-grade versus low-grade CAV are provided in **Supplemental Figure 2**.

We then turned our attention toward understanding T cell clonal dynamics in CAV. Using TRUST4^37^ to explore T cell clonal dynamics in CAV, we found no significant differences in the number of unique TCRs or TCR clonality across the spectrum of CAV grades (P-value > 0.05 for all comparisons by Kruskal-Wallis test; **Figure 3A** and **3B**). Similarly, there were no significant differences across a wide array of diversity indices between the CAV groups, though several metrics had P-values < 0.1 by Kruskal-Wallis test (Shannon index P-value = 0.06, inverse Simpson P-value = 0.06, entropy P-value = 0.06, Gini-Simpson index P-value = 0.08, and ACE P-value = 0.08; **Figure 3C**).

**Figure 3.**
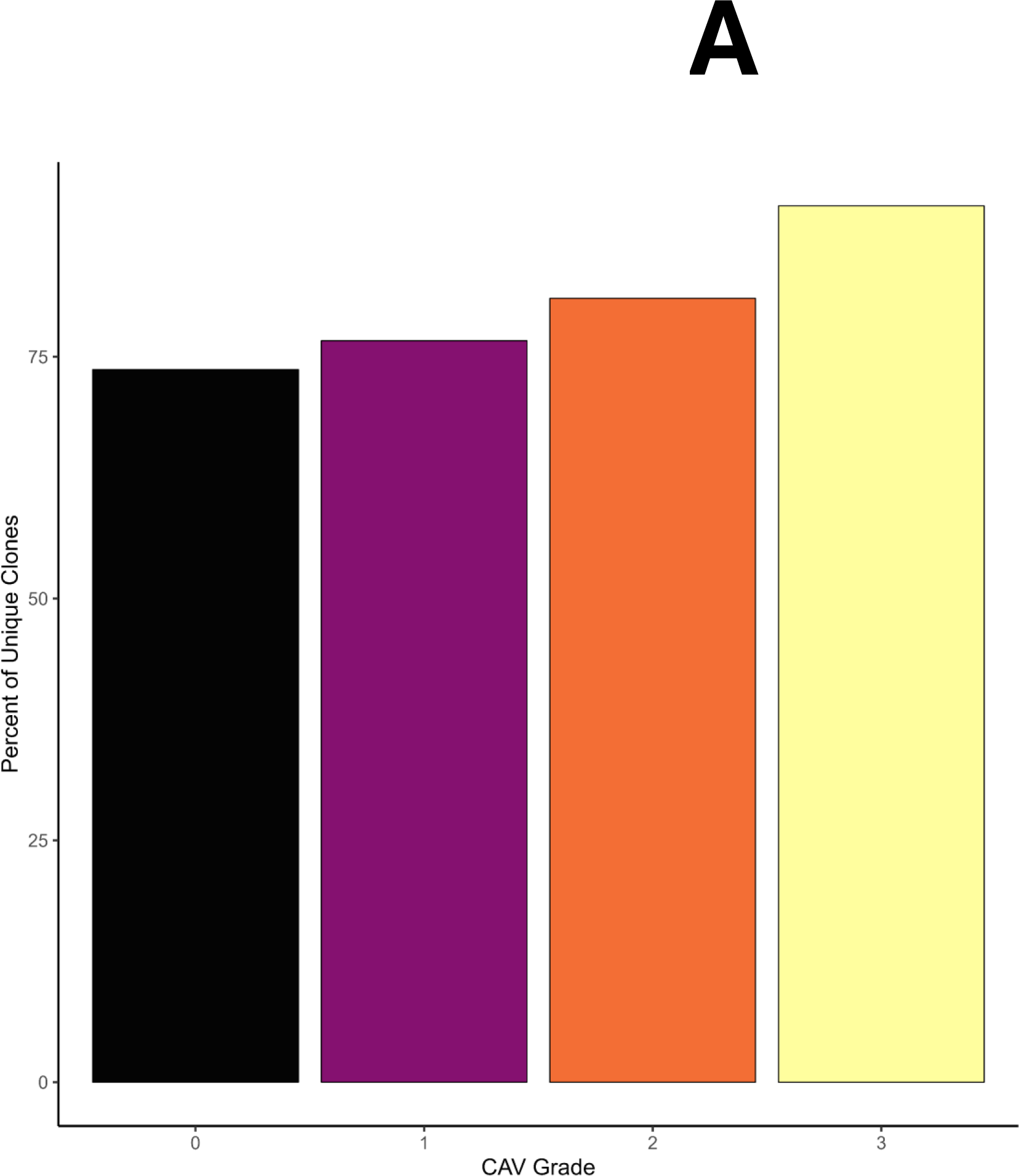
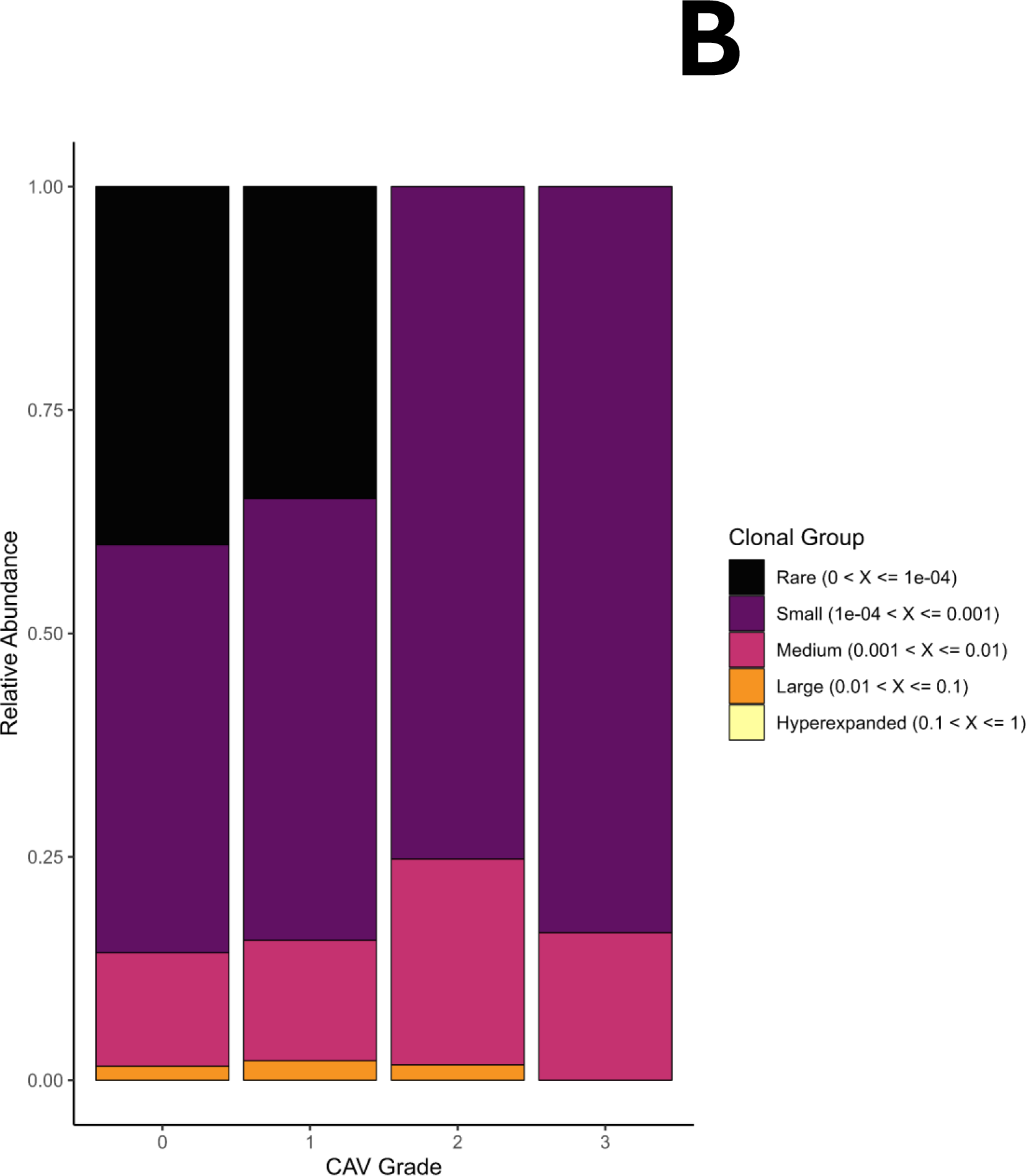
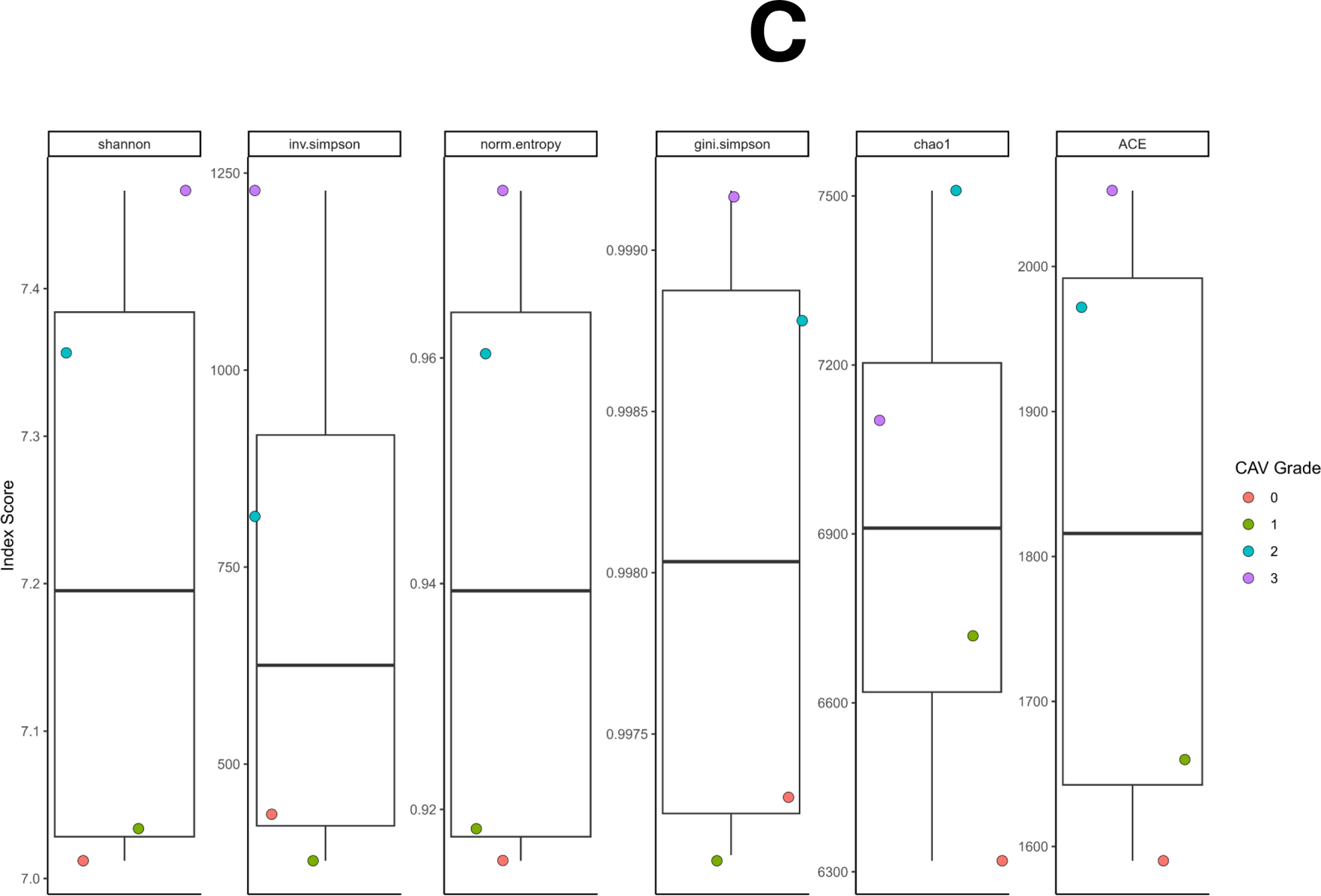
TCRs were reconstructed using TRUST4 and analyzed using scRepertoire. Using barcode matching with scRNA-seq, as described in the Methods section, ensured that TCR analyses only included T cells that passed rigorous quality control. **A)** Total number of unique TCR clones according to CAV grading did not show any significant differences between groups (Kruskal-Wallis P-value > 0.05). **B)** TCR clonality across the spectrum of CAV grading showed no significant differences across groups (Kruskal-Wallis P-value > 0.05). **C)** TCR diversity was measured by the following indices: Shannon, inverse Simpson, normalized entropy, Gini-Simpson, Chao1, and ACE. No significant differences in TCR diversity were observed amongst the groups (Kruskal-Wallis P-value > 0.05 for all indices).

## Discussion

To our knowledge, this is the first study to leverage single-cell multi-omics to comprehensively characterize cell-specific transcriptomic differences in HT recipients with and without high-grade CAV. We find that there are subtle, statistically significant increases in circulating CD4+ T central memory cells and CD14+ and CD16+ monocytes in high-grade CAV. Further, we demonstrate 745 unique genes that are differentially expressed in high-grade CAV and are enriched for putative pathways involving inflammation, angiogenesis and vascular smooth muscle cell proliferation, consistent with our current biological understanding of CAV pathogenesis. Importantly, these genes are differentially expressed despite adjustment for age, gender and prednisone use. Our findings suggest that the use of peripheral gene expression profiles may facilitate diagnosis of high- versus low-grade CAV, potentially obviating the need for invasive coronary angiography in the latter group.

Prior studies have shown mixed results in the use of mRNAs and miRNAs for detection of CAV^8, 10, 11, 20^. For example, the success of a targeted 11-gene panel (i.e., *Allomap*) for surveillance for acute cellular rejection has not translated consistently to CAV diagnosis^20^. The genes in this panel include those involved in glucocorticoid responses (*IL1R2*, *FLT3*, *ITGAM*), lymphocyte homing (*ITGA4*), immune checkpoints (*PDCD1*), and other non-specific pathways (*MARCH8*, *WDR40A*, *PF4*, *C6orf25*, *RHOU*, and *SEMA7A*)^6^. As CAV development is thought to be a more indolent process as compared to the acutely heightened inflammatory response that occurs in acute rejection, identification of changes in expression of only a small number of transcripts may be insufficient to characterize CAV. Furthermore, the *Allomap* assay is influenced by steroid use of >20 mg/day, transfusion products, and changes in immunosuppression regimens within the preceding 30 days, among other things^6^. On the other hand, whole transcriptomic analysis at cellular resolution allows for unbiased identification of a large myriad of genes that may be associated with high-grade CAV. In our study, following adjustment for age, gender and steroid use, we were able to identify 745 genes spanning across multiple cell types that are uniquely DE in high-grade versus low-grade CAV. That these DE genes show enrichment in putative pathways involved in CAV pathogenesis indicates biologic plausibility of our results. In contrast, miRNAs showed early promise in a number of diseases as potential biomarkers but have not been consistently reproducible even in the same disease^10, 11^. In part, this may be due to challenges with sample storage, miRNA isolation and quantification, and bioinformatics methods for analyzing miRNA data^54^. With mRNA, as in our study, there are robust pipelines for sample acquisition and storage, transcript quantification, and analyses^55^.

Others have attempted to measure serum cytokine levels or proteomics from different body fluids as a way to non-invasively diagnose CAV^12, 15–17^. However, serum cytokines are susceptible to significant fluctuations based on physiological changes, including diurnal variations, thus limiting their reproducibility even within the same individual over time^56, 57^. Proteomics approaches are primarily hampered by the need for targeted assays, which bias the results^21^. With scRNA-seq, we measure mRNA transcripts across the whole transcriptome, which minimizes the bias associated with targeted platforms, whether using genomic, transcriptomic, or proteomic assays.

While the strength of our study is in the interrogation of the whole transcriptome at single-cell resolution, this work does have important limitations. First, our sample size includes only 40 HT recipients, which precludes us from splitting the cohort into derivation and validation cohorts. Larger studies will be needed to validate our results. Second, we used novel statistical approaches for DE analyses^33^, which may be unfamiliar to some and require the use of scRNA-seq in order to identify small groups of cells that are perturbed in disease. These strategies (e.g., neighborhoods, metacells, etc.)^33, 58, 59^ increase sensitivity for detection of DE genes between conditions and are increasingly being used in the field of scRNA-seq. Finally, it remains unclear whether the cell-specific gene expression changes we identify precede development of angiographic CAV or are a manifestation of response to CAV. If the former is true, our findings offer the possibility of predicting future CAV development and progression, thereby identifying patients who may benefit from mammalian target of rapamycin inhibitor (mTORi) therapy.

Our results suggest that unbiased whole transcriptomic analyses at the single-cell resolution may identify transcriptomic-based biomarkers of CAV, offering the possibility of earlier and less invasive CAV detection in heart transplantation. Ongoing work is needed to explore longitudinal scRNA-sequencing of PBMCs during the course of CAV development to identify the timing of divergent cell-specific gene expression patterns between patients who go on to develop high grade CAV and those who do not.

## Author contribution

Conceptualization: K.A., K.H.S.; J.C.R.; Formal analysis: K.A.; Q.S.; Writing and revision: K.A., K.H.S., N.C., J.E.F; Funding acquisition: J.C.R.

## Supplemental Figures

**Supplemental Figure 1.** Single-cell WGCNA identified 8 modules of co-expressed genes in high-grade CAV, visualized here as **A)** beeswarm plot across cell types and **B)** overlaid on the UMAP. These modules represent genes listed **Supplemental Table 5**.

**Supplemental Figure 2.** Pathway enrichment analyses of module-specific genes in high-grade CAV. Shown are dot plots representing enriched pathways in module 4 (**A**), module 5 (**B**), and module 8 (**C**). The dot plot for pathway enrichment for genes in module 7 is shown in Figure 2D.

