## Supplementary Figures for "Single-cell RNA-sequencing identifies unique cell-specific gene expression profiles in high-grade cardiac allograft vasculopathy"

### Slide 1
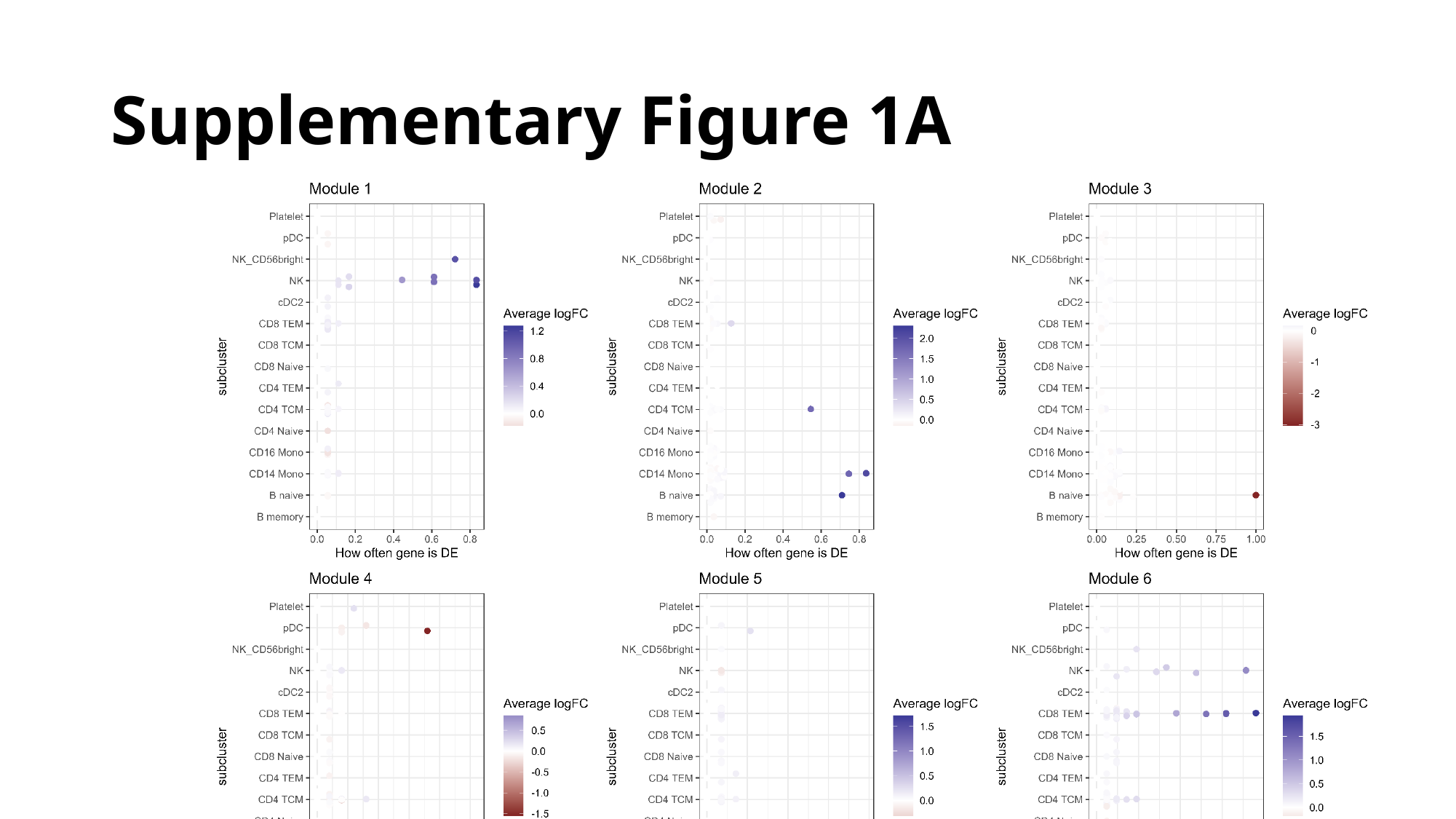

#### Supplementary Figure 1A

### Slide 2
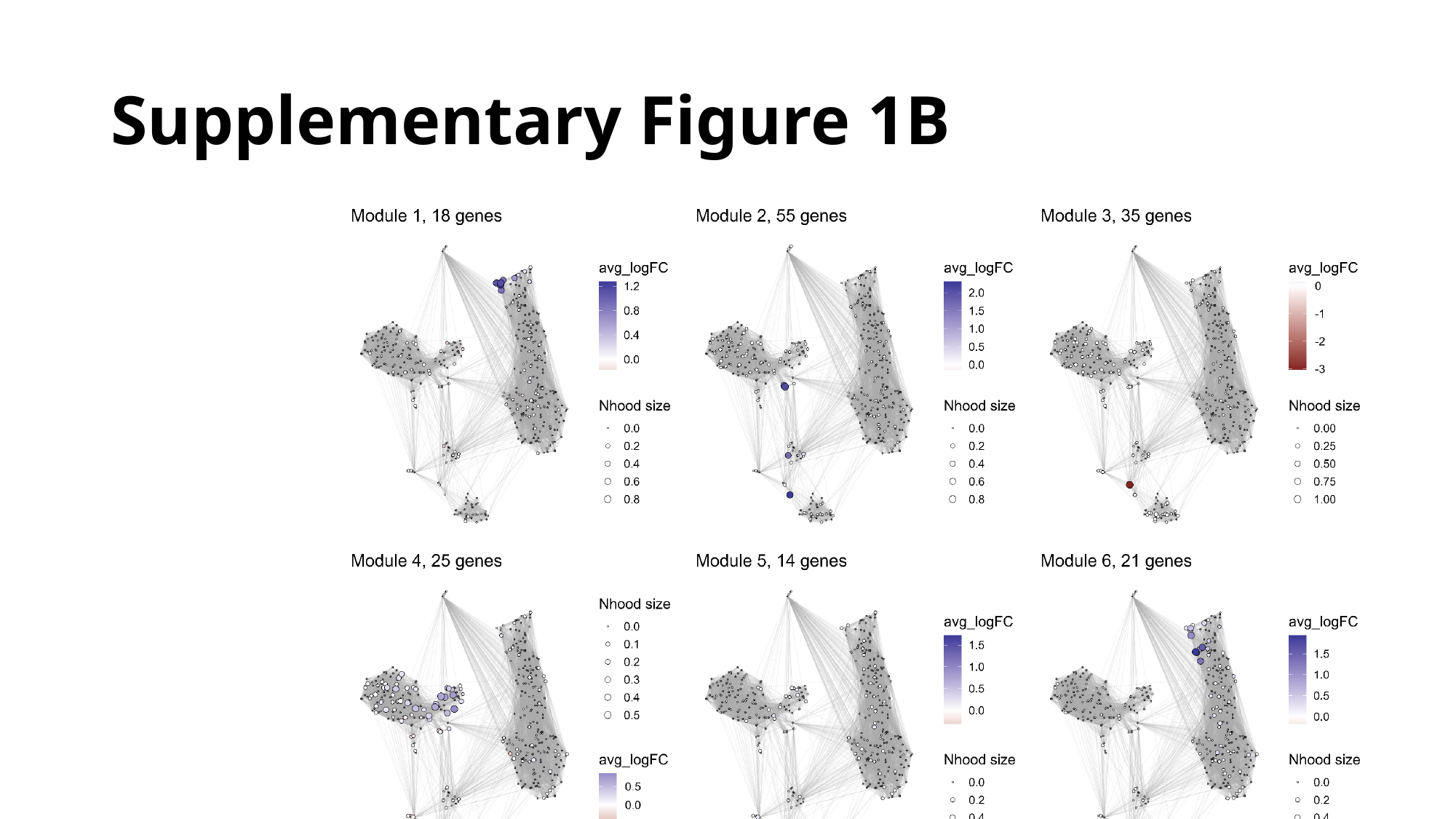

#### Supplementary Figure 1B

### Slide 3
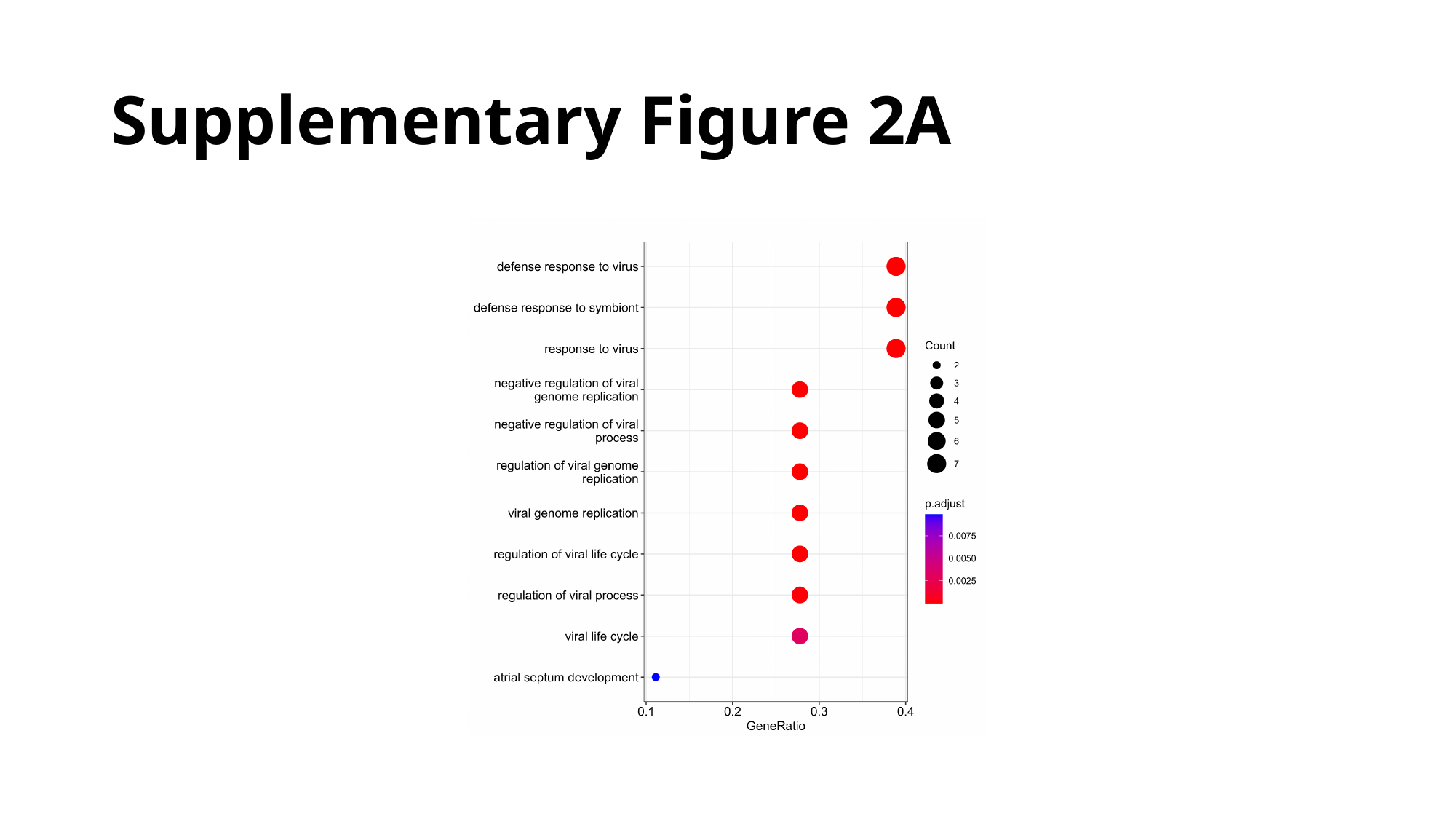

#### Supplementary Figure 2A

### Slide 4
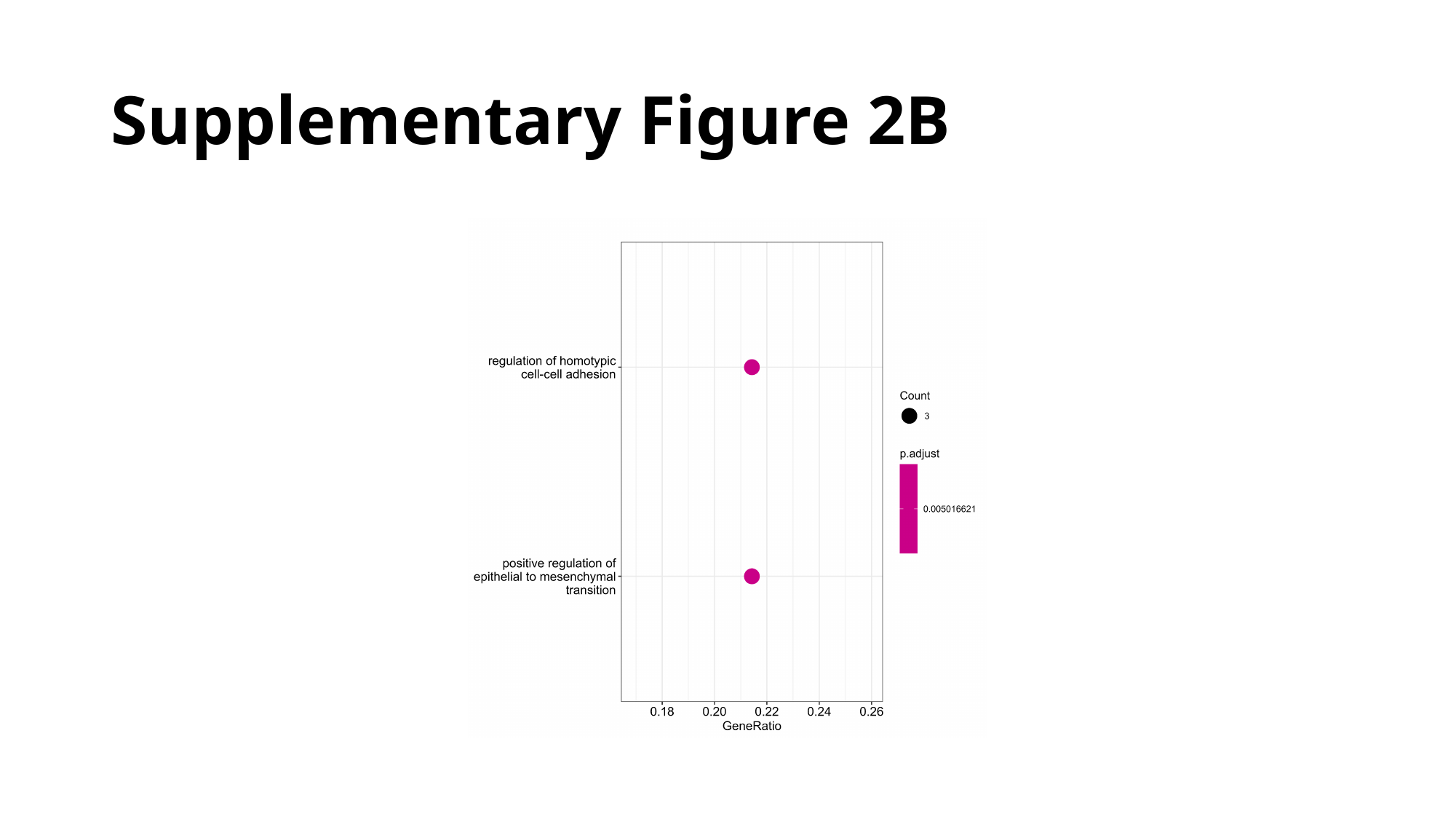

#### Supplementary Figure 2B

### Slide 5
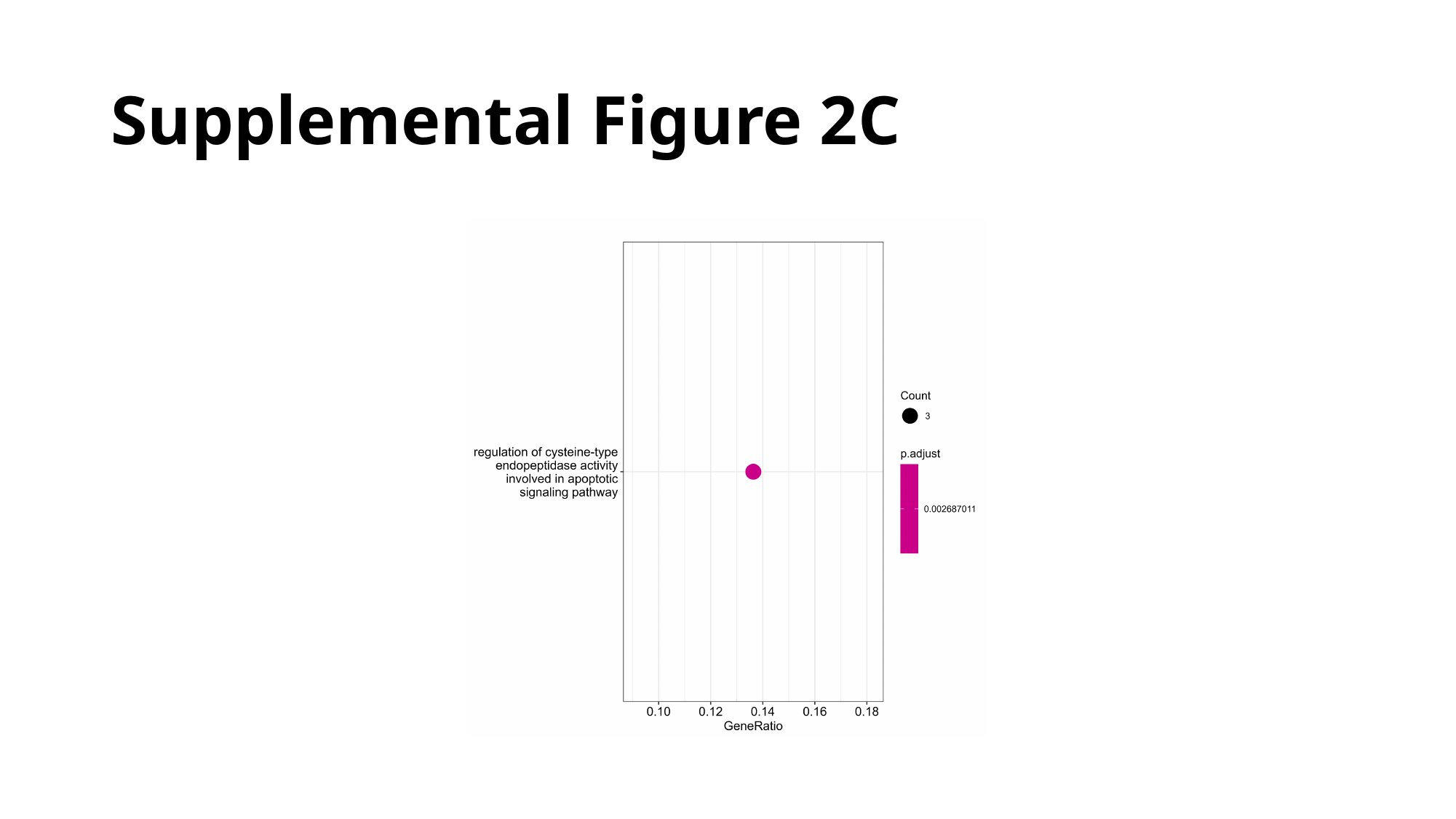

#### Supplemental Figure 2C
